# Effort distribution changes effector choice, behaviour and performance: A visuomotor tracking study using finger forces

**DOI:** 10.1101/230110

**Authors:** Satishchandra Salam, SKM Varadhan

## Abstract

Human movement and its associated performance are bounded by a hierarchy of constraints operating over certain control variables. One such variable of both physiological and behavioural importance is the mechanical effort exerted by the participating elements. Here, we explored how motor performance is affected by the distribution of work, and consequently the effort.

Using human hand as a model, we employed a visuomotor tracking task to study the associated motor performance when mechanical effort exerted by the fingers are modulated. The subject has to trace a set of ideal paths provided on visual feedback screen to reach a target through a cursor controlled by index and little finger forces. Modulation of these forces allows us to see how the perceived effort requirement affects the tracking performance. In this task demanding two-element coordination, we represent index finger as the independent/dominant element against little finger as the dependent/subjugate counterpart. We study how increasing mechanical effort contribution from the independent element leads to changes in both behaviour and performance.

We found that despite higher mechanical requirements of employing index finger to produce larger absolute force, the movement control system continues to prefer it as against little finger which could have produced smaller absolute force. Moreover, the observation of better tracking performance under larger contributions from the independent component reflects to a plausible hierarchy of constraints employed in the motor control system that operates with more than one objective, energy minimisation per se. At least for the behaviour in study, the improved motor performance suggests that the control system prefers higher independence of the participating elements.

## Introduction

The successful execution of meaningful and goal directed movement demands for the control and coordination of the participating elements. As it has been popularised by the Bernstein redundancy problem [Bernstein 1967], there are multiple equivalent motor solutions for the execution of a movement. This, in turn, facilitates variability of the movement — there are redundant or abundant [Latash 2012] ways of recruiting the required motor units for the execution of a movement. Yet with repeated movements and successful development of fitness solutions to the task requirements, patterns emerges (in the repeated movements) and it manifests itself into behaviour [Beer 2009], Together with, the study of this associated behaviour could elucidate the mechanisms of control and coordination involved in the generation of human movement.

In the context of this study, a variable of interest is the distribution of work, and subsequently the effort required, across the participating effectors. How does the motor control system recruit from the redundant set of effectors? Which properties of the effectors dictate the recruitment policies? It has been shown that a policy of minimising largely effort and marginally variability is adopted in an absolute finger force production task [O’Sullivan et al. 2009], A statistical decision theory outlook speculates that these choices could be determined by the associated gain and loss functions [reviewed in Wolpert et al. 2012], Or for the generation of movement trajectories in spatial space, various cost functions have been suggested including minimum jerk principle [Flash et al. 1985] and minimum intervention principle [Todorov et al. 2002], Following the theory of signal dependent noise, the associated variability due to the ‘noise’ in the motor command should increase with increase in the size of the control signal itself [Harris et al. 1998], Further, such models that also accounts for the effort cost function (along with a few other constraints) have simulated qualitatively similar movements [Guigon et al. 2007].

Thus, given how the motor behaviour and performance is influenced by the participating elements, the choice of effectors could be influenced by how the effort distribution across the participating effectors yields to changes in motor performance. In this experiment, we used a visuomotor tracking task which demands production of dynamic and precision finger force (for the successful execution of the task) to study the associated changes in behaviour and performance. By modulating the visual feedback across different effort requirements for the execution of the task, we study the effects of relative mechanical effort contribution on effector biasing, tracking accuracy, control and speed.

Particularly, for this task of visuomotor tracking using finger forces, the sensory information which could primarily affect the optimal performance are derived from vision, cutaneous receptors and proprioception. Through studies on intermittent force production using visual feedback, the role of vision in estimating the ‘missing’ information have been established [Miall et al. 1993, Slifkin et al. 2000], The touch of the fingertips on the sensor provides an interface to give somatosensory feedback to the motor control system which contributes towards optimal performance of the task. This is partly due to the cutaneous receptors present on the hand whose role have been established through studies of grasping and object manipulation [Johansson et al. 1984, 1992], The other source of somatosensory information is the proprioceptive information which can be accessed from the involving motor units [Matthews 1964], Patient evaluation has also clarified the deficits in motor functionality with impaired proprioception [Rothwell et al., 1982, Sanes et al. 1984], And lastly from temporal perspective, across a wide and inconclusive estimations, the temporal capacity of the short visuomotor memory for the task involving finger force production through visual feedback is estimated to be around 0.5 s - 1.5 s [Vaillancourt et al. 2002].

Following the concepts of enslavement [Zatsiorsky et al. 1998] and spillover [updated review in van Duinen et al. 2011], we used index and little finger to represent independent and less-independent pair, or independent-dependent pair (for nomenclature purpose in this binary coordination task). For the lack of definition, analogies are drawn for this pair into as dominant-subjugate pair, and also as stronger-weaker pair. The results showed that the motor control system has a preference for using the more independent effector compared as against its counterpart. This behaviour manifests into improvement of tracking accuracy and control with increasing contribution of relative mechanical effort from the independent element. These results provide insights about how the movement control system realises certain perceived and performing behavioural parameters. It has critical implications in how the control and coordination is achieved in the redundant multi-effector system. In addition, this study introduces a potential behavioural method to measure the relative neural biasing acting upon the pair of participating elements.

## Methods

### Participants

10 subjects (5 males; age: 25.20 ± 3.29 years, mean ± standard deviation) from the student population of Indian Institute of Technology Madras (IITM), India, were recruited for the experiment. All the subjects reported being right handed according to their use of writing, and had no history of any neuromuscular disorders which could interfere with the pressing tasks. Only the explanation of the experimental tasks was provided to the subject, and they were naive to the purpose of the experiment. Also, a monetary reward of INR 500 was provided at the successful completion of the session. They read and signed an informed consent document. All experimental procedures were approved by the Institutional Ethics Committee of IITM, India (IEC/2016/02/VSK-7/17).

### Experimental Setup and Data Acquisition

Two force sensors (Nano-17, ATI Industrial Automation, USA) capable of measuring force and torque in all orthogonal three axes and three planes (respectively) were used for measuring the index and little finger forces. To prevent the slippage of fingers over the sensor surface, and to reduce possible physical environmental contamination (such as humidity), sandpaper of grit size 100 was used to cover the sensor surface. The sensors were fitted on a platform with slots to facilitate the adjustment of sensor position to finger lengths of different subjects. The finger forces were sampled at 200 Hz. A customised LabVIEW environment (LabVIEW 2014, National Instruments, Austin, TX) was used to interface and provide the visual feedback through a 21 inch screen placed 0.75 m in front of the subject.

### Tasks

The experimental tasks consisted of three different subtasks: 1. Maximum force production task, 2. Constant force production task, and 3. Tracking task.

#### 1. Maximum voluntary contraction task

In this task, the normal component of maximum (isometric) voluntary contraction (MVC) force of the individual fingers (index: I and little: L) were measured. A visual feedback for the time profile of the normal component of the finger force was provided on the screen. Subjects were instructed to produce their maximum finger contraction force within a 10 second duration. Trials were repeated for 3 times with a 1 minute interval in between. The highest value were taken as reference for the following tasks. A 3 minute break was given at the end of the task to avoid any possible development of fatigue.

#### 2. Constant force production task

Through the visual feedback (*Figure Set-up*) provided on screen, subjects controlled a cursor using index finger force along horizontal axis and little finger force along vertical axis. In this task, the subject has to bring and hover continuously the cursor over the target positions as accurately as possible for a 15 second duration. (Pilot studies showed that subjects were capable of performing the navigation task successfully in about 10-20 second.) The targets represent 15% of MVC for index finger, and 15, 10, 7.5 and 5 % MVC for little finger. Also, inter-trial breaks of 30 second were provided between.

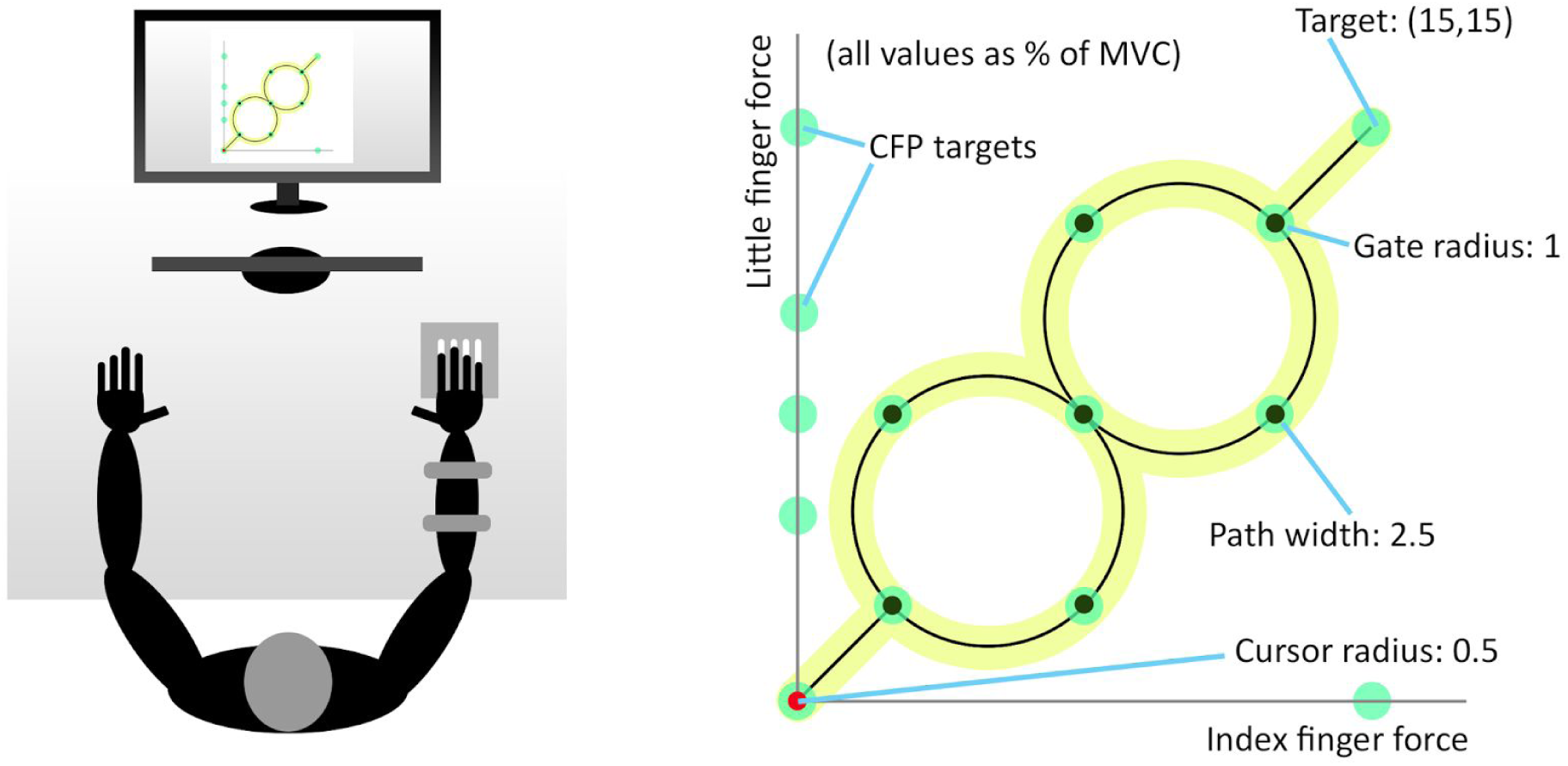
Figure Set-up: Experimental setup (left) and visual feedback (right). Nodes and gates are included along the ideal paths to assert the choice of a path. The targets on the axes are for the constant force production task.

#### 3. Tracking task

The visual feedback screen shows a redundant set of ideal paths consisting of two straight line segments and two visually perfect circles. A target point representing specific finger forces combination was marked at the outer end of the path. The subjects were instructed to “reach the target about any of the ideal path”. The cursor which has a finger force proportional displacement has to track about any of the ideal paths to reach the target. This requires that the subject has to produce specific combinations of force to navigate around and trace about the ideal paths to reach the target. The associated motor behaviour was investigated across relative mechanical effort (that should be exerted by the participating elements) expressed through:

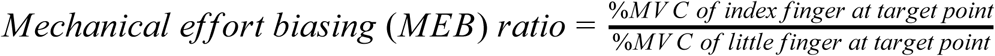

For this experiment, the mechanical effort is computed as the MVC-normalised-force produced by a finger, i.e., it is the relative amount of force generated by a finger with respect to its MVC. Four different experimental blocks were conducted on four values of mechanical effort biasing (MEB) variable defined as the ratio of mechanical effort of index finger force to the mechanical effort of little finger force at the target point. Hence, the final target corresponds to 15 - 15, 15 - 10, 15 - 7.5 and 15 - 5 % MVC forces of index and little finger respectively for corresponding MEB ratios of 1:1, 1.5:1, 2:1 and 3:1.

Explanations were provided to maintain a practical accuracy implying that they don’t do any unusual actions such as moving the cursor either extremely too slow or too fast. This was done to achieve a practically consistent set of performance across the subjects. Each trial was started when the subject responded his/her readiness at the audio cue provided by the experimenter. In addition to the breaks provided anytime at the demand of the subject, a 3 minute break was provided at the end of each block.

### Experimental protocol

The subjects performed the constant finger force production using the MVC recorded in the preceding task (*Figure Experimental Protocol*). For the navigation task, it requires that the subject continuously produces a dynamic and unique combination of finger forces within a permissible range of error. Such a task posits a higher motor skill requiring individual’s unique ability to perform; and hence following the saturation of skill acquistion in the motor learning paradigm, a training session was provided for the subject at the beginning of each block to learn and acquaint with the novel visuomotor task. Only when the subjects were capable of performing ‘good’* in the training session (lasting about 10 - 20 trials), they proceeded to conduct the navigation task. It is to assume that the subjects have reached the ‘saturation’ level in the training-performance curve. Of 10 subjects recruited for the main experiment and 5 subjects for the pilot experiment, only 1 subject was unable to complete the training successfully.

*Evaluation of a trial was done largely through online observation of the performance by the experimenter. As the training progresses, the subject exhibited visually acknowlegeable improvement and saturation of tracking performance. At the end of about 20 minute of free training to the novel task, an online statistic called stay percent was used to qualitatively judge the tracking performance. The stay percent measures for how much the cursor stays inside the 2.5% MVC wide path. A consistent performance across 5 consecutive trials above approximately 70% stay percent was considered sufficient to successfully finish the training.

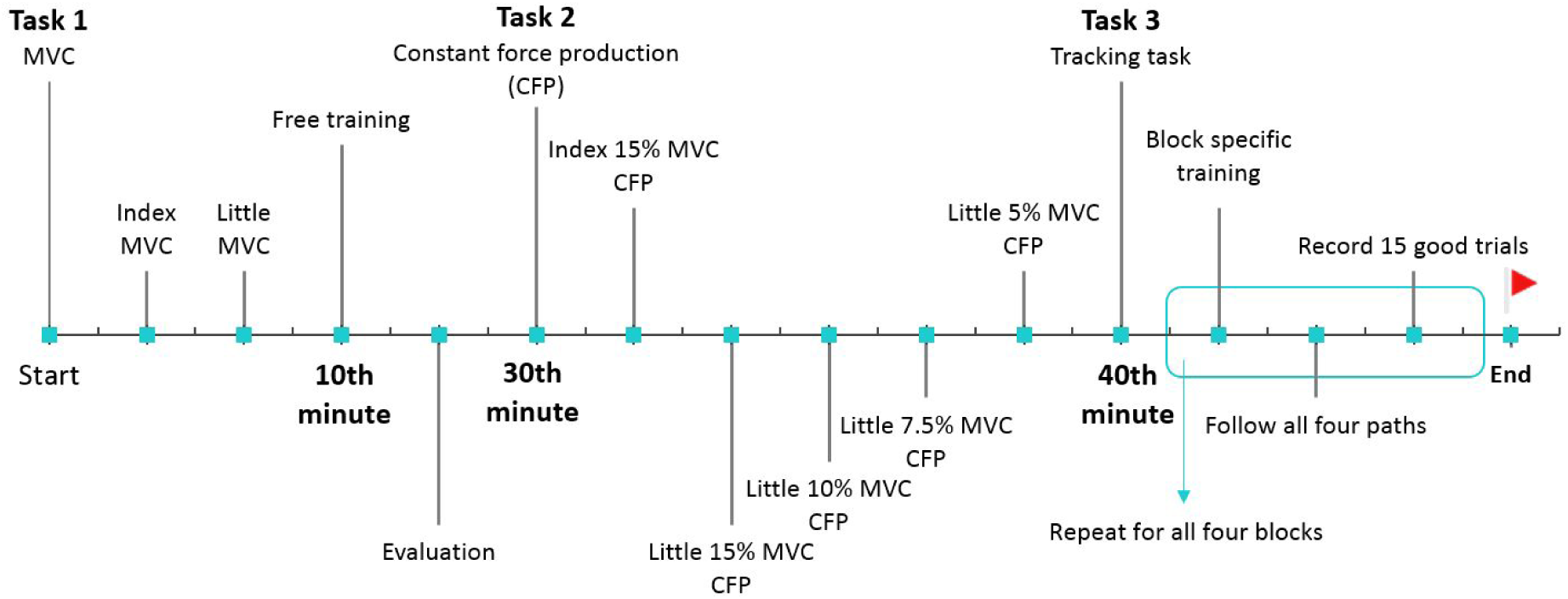
Figure Experimental Protocol: Before the tracking task in Task 3, Task 1 normalises the effort requirements across different subjects with different abilities.

### Procedure

The subject seated comfortably on a height adjustable chair with their forearms rested on the table (Figure *Set-up*). Velcro straps were used to constrain the movement of the forearm during the experiment. The sensors were placed directly below the right hand of the subject where the subjects could press onto the sensors comfortably while looking at the visual feedback screen. The task specific instruction was provided at the beginning of each task. A typical navigation tasks lasted for 15 - 20 second. Including the breaks between the sets, the whole set of tasks were completed in about 1 hour 20 minutes.

### Data analysis

The finger force data were digitally smoothed using a fourth-order zero-lag Butterworth filter with a cutoff frequency of 15 Hz. Four sample trajectories of a cursor following about the ideal path across the four experimental blocks are shown in *Figure Sample trajectories*. As it can be seen, the mechanical effort generated by the index figure is fixed at 15 % MVC, while the mechanical effort of little finger changes across different experimental blocks.

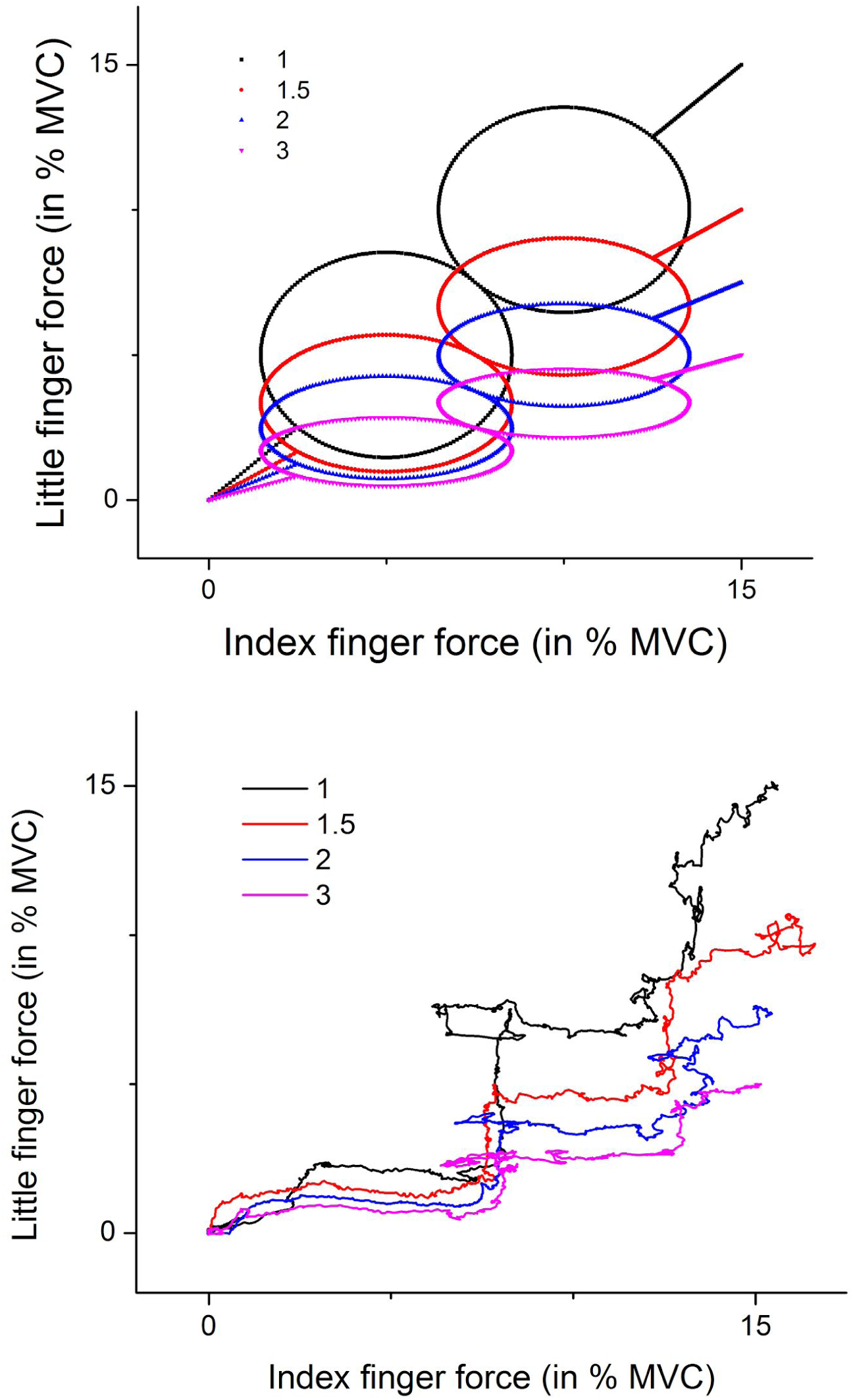
Figure Sample trajectories: (Top) Ideal paths in force space. Subjects can follow any of the four ideal paths. (Bottom) Representative sample trajectories across 4 blocks of MEB ratio. While the visual feedback remains the same across all four blocks, the representations in the force space changes across blocks. Effort contributions by little finger changes across blocks. The final target corresponds to (15,15), (15,10), (15,7.5) and (15,5) %MVC of (index, little) finger.

### Visual and force space

The visuomotor task in this experiment is built on the kinetic space of the finger forces. The cursor provided on the visual feedback screen has a force proportional displacement of the index finger force along the horizontal axis, and the little finger force along the vertical axis. As the feedback is modulated across different experimental blocks with change in mechanical effort biasing ratio, two distinct spaces emerges in this visuomotor task: the visual space - as it is seen in the feedback screen; and the force space - as what amount of force has actually been produced. Hence, two distinct statistics of tracking performance on both the spaces are calculated for the same trial.

### Tracking error

During the course of trajectory, tracking error at any instant is calculated as the minimum Euclidean distance of the trajectory point (at that instant) from any of the ideal path (*Figure Tracking error*). Further, directionality is assigned to represent the biasing of the cursor towards either index(+) or little finger(-). The visual tracking error is calculated by first transforming the force values into as what is appeared on the visual feedback screen, i.e., into slope-one straight line segments, and perfect circles. On the contrary, the force tracking error is calculated by transforming the ideal path into the transformed ideal path, i.e., into slanted straight line segments, and vertically compressed ellipses.

For testing the normality of the series, Anderson-Darling test was done by using MATLAB function ‘adtest’ from Statistics and Machine Learning Toolbox.

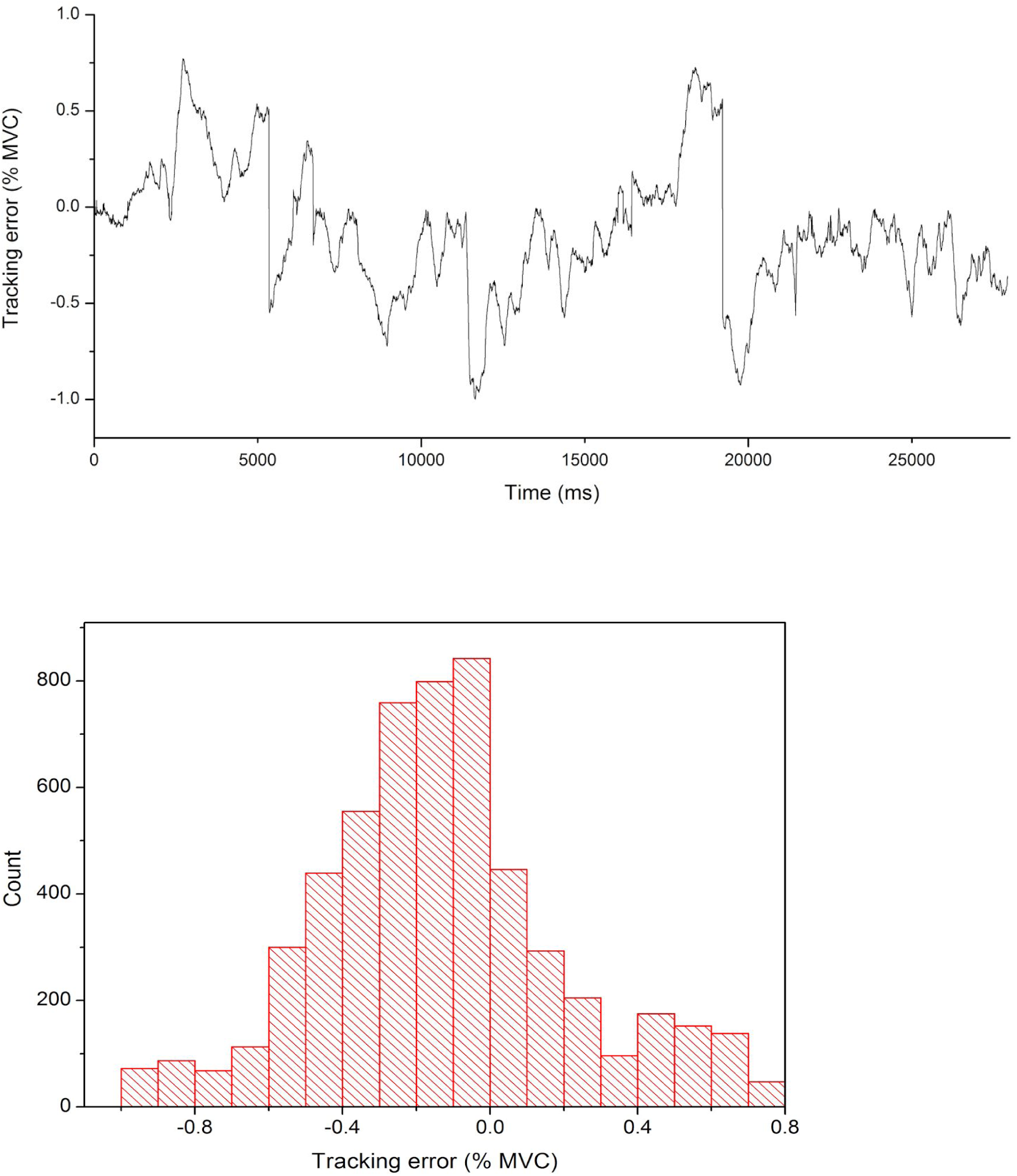
Figure Tracking error: Top: Sample tracking error series of cursor about the ideal path from block of MEB ratio 1:1. The series is the same in both force and visual space for MEB of 1:1. Bottom: Histogram of the tracking error series.

### Biasing of trajectories

This biasing of a trajectory of a trial is computed by calculating the mean of the tracking error series. Following the sign convention adopted earlier, a negative mean corresponds to index finger biased trajectory and a positive mean to little finger biased trajectory (*Figure Biasing map*).

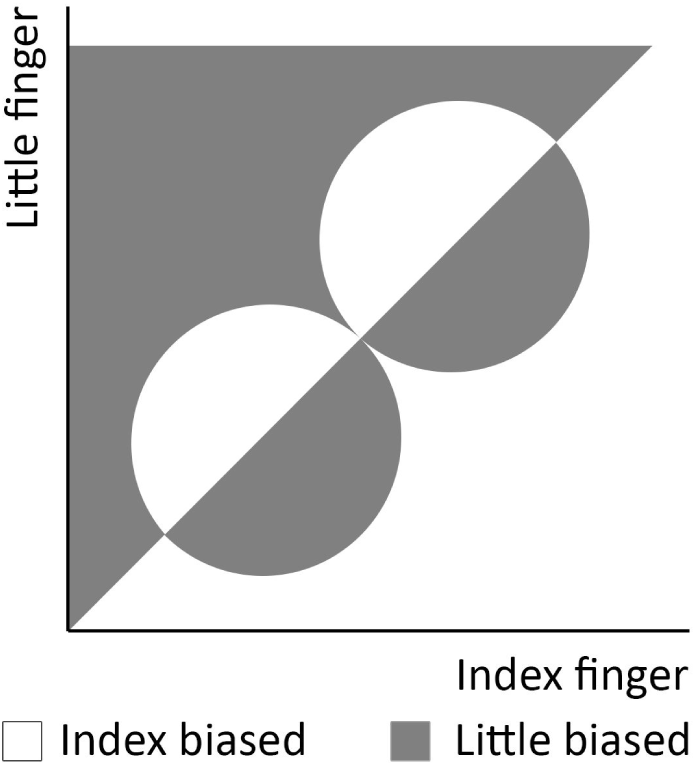
Figure Biasing map: Within the operating space till (15,15) %MVC of both fingers in visual space, shaded areas represent little finger biased trajectory points and the unshaded area represents index finger biased trajectory.

### Interaction correction of biasing

For the involved pair of effectors, since it belongs to the same control system, they need not be purely independent and may interact. This interaction is incorporated into the biasing result by modifying the performed trajectories into space which accounts for the interaction. The ideal and performance trajectories are transformed with interaction coefficients - coefficients which represents the unintended production of force when the other effector is in action.

As mentioned in Task 1, the MVC was recorded while providing a visual feedback of temporal profile of the finger force and without explicit instruction to follow any systematically increasing force profile. This renders the estimation of interaction coefficients from the dataset analytically complicated. Thus, for this paradigm using finger forces, enslavement coefficients from Zatsiorsky et al. 1998 are used to correct the observed biasing result (*Table Interaction coefficients*). Further, it has been assumed that the interaction coefficient doesn’t change with change in effort.

Table Interaction coefficients: Only concerned values involving index and little finger are shown (in %MVC units). Adapted from Zatsiorsky et al. 1998.

| Master (column) | I | L | I/L (symmetricity) |
| --- | --- | --- | --- |
| I | 100 | 10.9 | 1.31 |
| L | 14.3 | 100 |  |
| IL | 79.3 | 75 | 1.06 |
| IMRL | 67 | 63.8 | 1.05 |

Statistics of tracking performance: The performance of a trial is evaluated along two orthogonal dimensions: the performance (1) about, and (2) along the ideal path. The performance about the ideal path measures the sidewise sways about the ideal path. And the performance along the ideal path measures the forward and backward progress that the cursor makes during the course of the trajectory.

### Performance about ideal path

The visual variability (vVar) measures the performance as it is appeared on the visual feedback screen. It measures the deviation as it is exactly seen in the screen. On the other hand, the force variability (fVAr) measures the kinetic performance. It measures the deviation of the actually generated force from what should have been generated to trace the ideal path. Once again, following the approximately normally distributed tracking error series, its root mean square is considered as performance variability (as a statistic of motor performance). And the inverse of this variability is interpreted as the motor accuracy (stictly, precision). These performance statistics were averaged across the 15 trials for the 4 mechanical effort biasing (MEB) variables for all 10 subjects.

### Performance along ideal path

For a system which has a ‘good’ control over the end effector, the trace of the cursor would be a cumulative series of trajectory points which makes forward progress only. The cursor going backwards instead at any point is an indication of ‘poor’ or ‘loss’ of control. In the trajectories traced by the cursor in this visuomotor task, the control that the system has over the cursor is poor enough to make considerable amount of backward corrections. Here, the ratio, called the correction ratio, of the forward progression to the backward movement is used to measure this performance of trial along the ideal path. It is (similarly) averaged across the 10 subjects, and the corresponding error of mean is also calculated.

### Speed of a trial

The average speed of the trajectory represents the rate of change of finger forces. It is computed as the distance traversed by the trajectory by its trial completion duration. Even though the trial completion duration is same in both the force and visual space, the distance traversed in the visual space and the force space are not the same (*Table Distance traversed*). Thus, similar to the variability indices, the average rate of change of finger forces are calculated in both the spaces.

Table Distance traversed: The actual distance traversed are slightly greater than the ideal distance. Distances are as mean±standard error for 10 subjects in units of %MVC.

| **MEB ratio** | **1** | **1.5** | **2** | **3** |
| --- | --- | --- | --- | --- |
| Ideal | 36.42 | 31.43 | 29.22 | 27.28 |
| Actual | 48.14±3.34 | 31.03±1.55 | 28.26±1.33 | 29.90±2.48 |

All these representative statistics for a trial are then averaged for the 15 trials for the 10 subjects across the 4 blocks of MEB ratio.

## Results & Discussion

### Normality of the tracking error

For all 600 tracking error series (15 trials, 4 blocks, 10 subjects), the Anderson-Darling test returns true for the normality with 95% confidence bounds. This consolidates the statistical basis of using mean and root mean square as estimators of the performance statistics of the 15 trials for 10 subjects across 4 blocks.

### Biasing in the two-effector system

For this task of tracking a set of paths in the force-force space of the finger forces, it is required that the participating effectors contribute their corresponding specific mechanical effort to be at a particular point across the course of the trajectory. And for the effectors involved, through concepts of enslavement [Zatsiorsky et al. 1998] and spillover [van Duinen et al. 2011], it has been established that index finger is the more independent finger as compared against the little finger. Further, drawing analogies with the effectors involved in this paradigm, index finger represents the independent-dominant-strong element with respect to the little finger as the dependent-subjugate-weak element.

The mean of the tracking error series is used to represent the biasing of the control system towards any of the participating elements. The result show that the trajectories thus generated are inclined towards the index finger (*Figure Biasing*). With increasing MEB ratio, that is, with relatively increasing mechanical effort contribution from the index finger with respect to little finger, the biasing of the effectors dissolves. A phenomena resembling a compensation or trade-off between effort and performance takes place; only at about 3:1 MEB ratio (15% MVC index, 5% MVC little), the biasing ratio tends to zero, which should correspond to unbiased control. This is a manifestation of the index finger producing more than the ideally required force thus resulting into the ‘pull’ of the trajectory towards the index finger axis. It implies that the control system has a preference of using the more independent effector compared against its counterpart.

For the pair of effectors chosen in this paradigm, owing to its neuromotor architecture, they interact with each other and interferes with their individual output. The production of force by the index finger will lead to unintended production of force in the little finger, and vice versa [Danion et al. 2003], This implies that they are not exactly an independent pair of effectors and this could influence the observed biasing result. The compensation could be made by correcting the actual trajectories to accommodate the interaction effects. For this paradigm using finger forces, this interaction could be quantified using the enslavement coefficients (with certain assumptions such as effort independent interaction). Curtly, since mostly symmetric interactions exist between the involved fingers [Zatsiorsky et al. 1998], the biasing result thus reported here should not be changed much even after the correction — which was observed in the result (*Figure Biasing*).

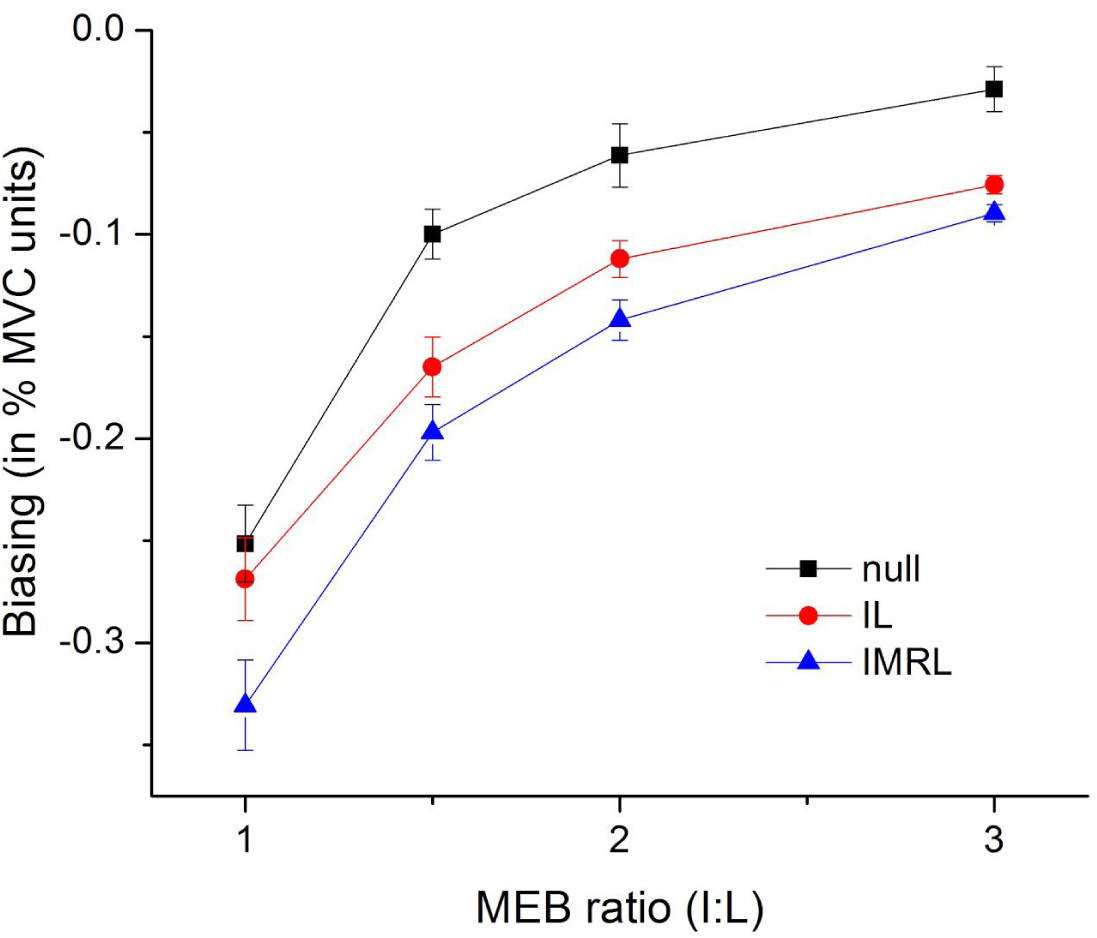
Figure Biasing: -ve for index bias, and +ve for little finger bias. In consequence to not instructing any explicit finger configuration, different subjects placed their IL (only I and L), IMRL (all fingers) or combination of both on the sensors. Thus the correction are shown for these two modes. ‘Null’ corresponds to no correction.

### Variability - performance about ideal path

The design of this experiment yields motor performance in two distinct spaces: force space and visual space. Hence, the motor variability (as a measure of motor performance) are computed in both these spaces (*Figure Variability result*). All statistics of variability decreases gradually with increasing MEB ratio. Also, the rate of drop of force variability (fVar) is higher than the rate of drop of visual variability (War). Hence, for this set of fingers (index and little) and for the mechanical effort range (within 15 % MVC both fingers), the performance (inferred as reduced variability) increases with increasing MEB ratio.

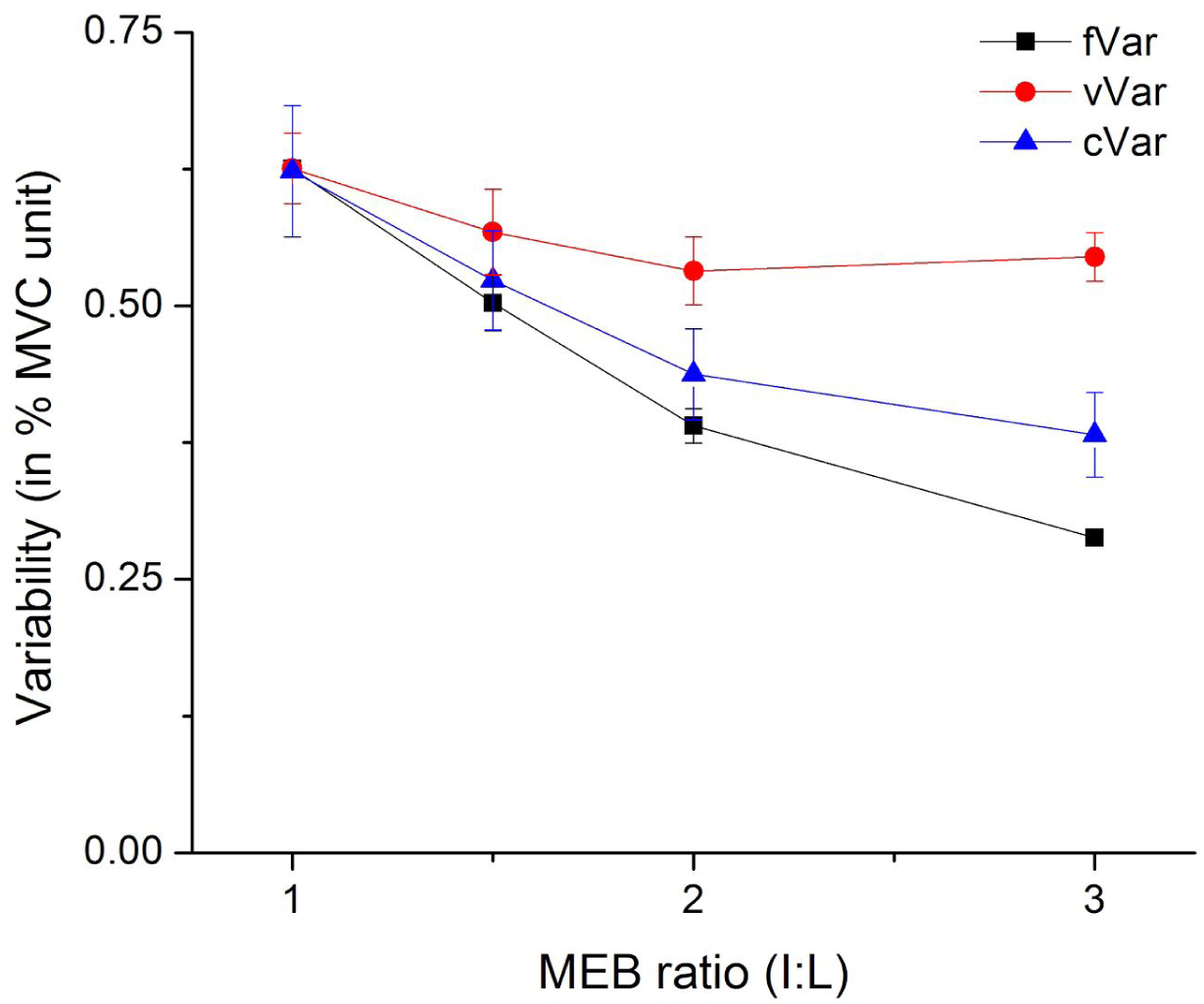
Figure Variability result: The root mean square of the tracking error series is used to represent the tracking performance variability; Variability as fVar in force space, vVar in visual space, and cVar in constant finger force production.

### Correction ratio - performance along ideal path

Ideal trajectory for a cursor to reach a target from a starting point would be a straight line connecting the two points. But as in this case of trajectory generated by two finger force production, the quality of the control is poor. Such imperfect performance resulting to the forward and backward sways of the cursor along the trajectory is quantified here.

The correction ratio, calculated as the ratio of forward progress to backward progress within a trial, increases with increasing MEB ratio (*Figure Correction ratio.*). There is a large distribution of this performance index across subjects (and hence the larger SE), and yet the pattern remains the same. This index also shows that the performance initially increases and saturates with increasing MEB ratio, as it was similarly observed with the variability statistics. In addition to the improvement in motor precision with increasing MEB ratio (from variability result), the increase in correction ratio also marks the improvement of motor performance in the sense that more forward movement are being made relative to backward movement.

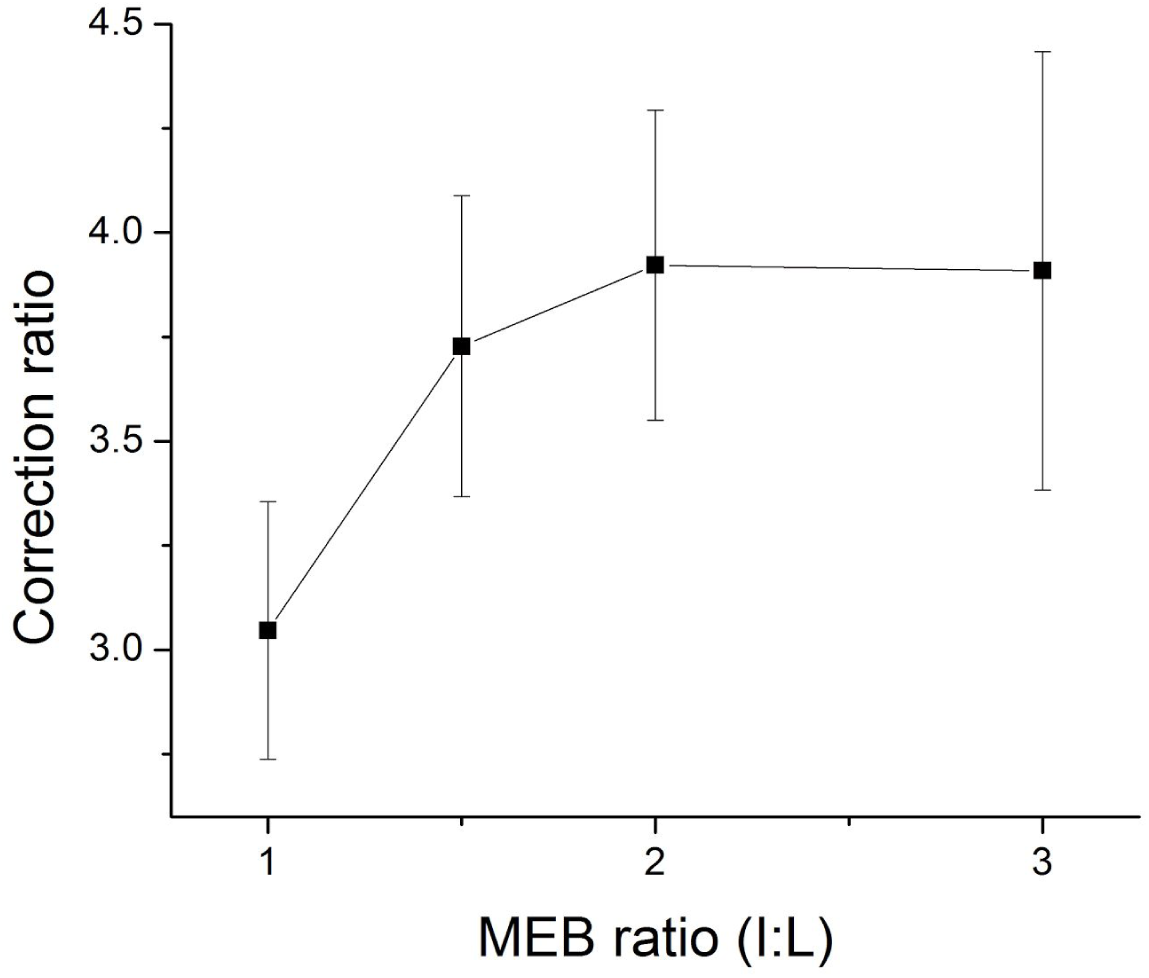
Figure Correction ratio: It also shows that the associated motor performance improves with increasing effort contribution from index finger — in the sense that the control gets better, and lesser backward movements are made.

### Average speed and speed-accuracy trade-off

Similar to the calculation of performance statistics in both the force and visual spaces, the average speed of the trajectory is also calculated for both these spaces (*Figure Average speed*). The result shows that the average speed of tracking decreases with increasing MEB ratio, which is the opposite trend of what was observed in the tracking accuracy. If all the performance variables associated in this paradigm were to improved with increasing MEB ratio, then the average tracking speed should also increase. Unlike what was marked as an improved motor performance in the tracking accuracy with increasing MEB ratio, this decrease in tracking speed is actually an indication of decline in absolute motor performance.

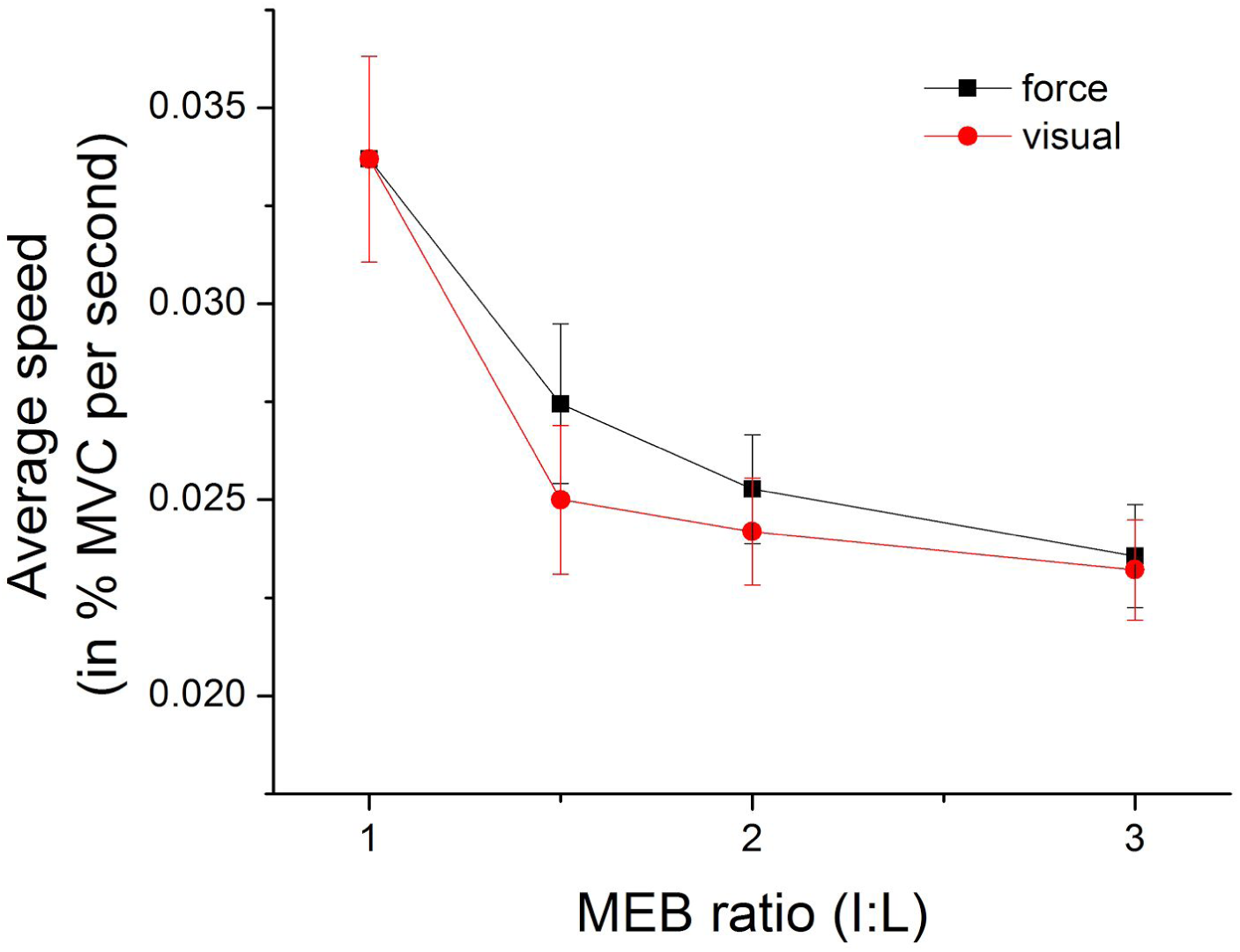
Figure Average speed: Despite the decrease in ideal distance to be travelled in force space, average speed decreases with increasing MEB ratio.

These contradicting observations could be due to multiple constraints operating over the control system. One such constraint could be the trade-off between speed and accuracy [Fitts et al. 1964] as it has been popularly established in task in kinematic space. But do similar principles of speed-accuracy trade-off in the kinematic performance apply to the kinetic performance variables? This could be supported by the fundamental mechanism through which human movement is generated. Movements are manifestations of the force generated by the participating elements and it is highly plausible that such similar trade-off policies applies to the kinetic performance variables as well.

In addition to this is how the rate of finger force is largely a task irrelevant parameter (*Figure Autocorrelation function*). This could mean that the decrease in the tracking speed is not due to the control system tracking slowly; this is what is resulted through the control of other variables - the control system could care less about the tracking speed. This is in conformation to the task instruction which does not provide any explicit instruction on the tracking speed.

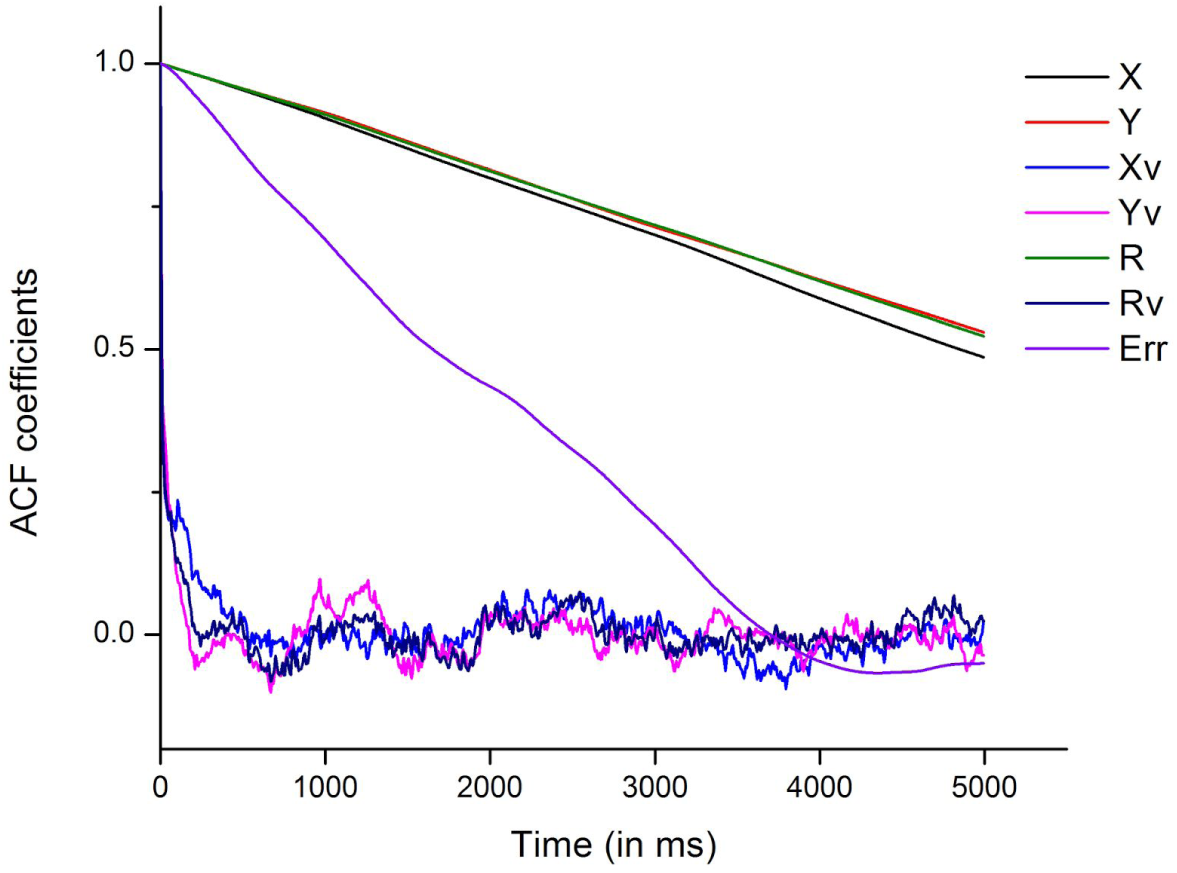
Figure Autocorrelation function: ACF coefficients across lags up to 5 seconds for a representative trial. ACF coefficients of rate of change of finger forces having a small value immediately beyond short lags implies that they are task variables of low task relevance [van Beers et al. 2013], X: index force; Y: little force; Xv: rate of change of X; Yv: rate of change of Y; R: position vector of trajectory point; Rv: rate of change of R; Err: tracking error.

In addition, the speed vs accuracy shows an inverse relationship; trade-off relationship do exists at least (*Figure Speed vs accuracy*). For these cloud of points, there are two possibilities: either (1) they belong to the same function, or (2) they belong to different effort-specific functions. For the first case, effort distribution would not affect the observed cloud of points; they all would have belong to the same function. But for the second case, as how true skill acquisition should be reflected on a systematic change in the speed-accuracy function [Reis et al., 2009; Shmuelof et al., 2012], a shift in the trade-off function should be observed with change in effort contribution. The cloud of points should belong to effort specific functions. But due to lack of any computationally established function supported by theories of motor control which could be used as a basis to fit over these points, it cannot be established whether which of these cases is true. Further experiments with speed and/or accuracy constrained conditions on the similar paradigm should elucidate the role of effort distribution in the shift of the speed-accuracy trade-off function.

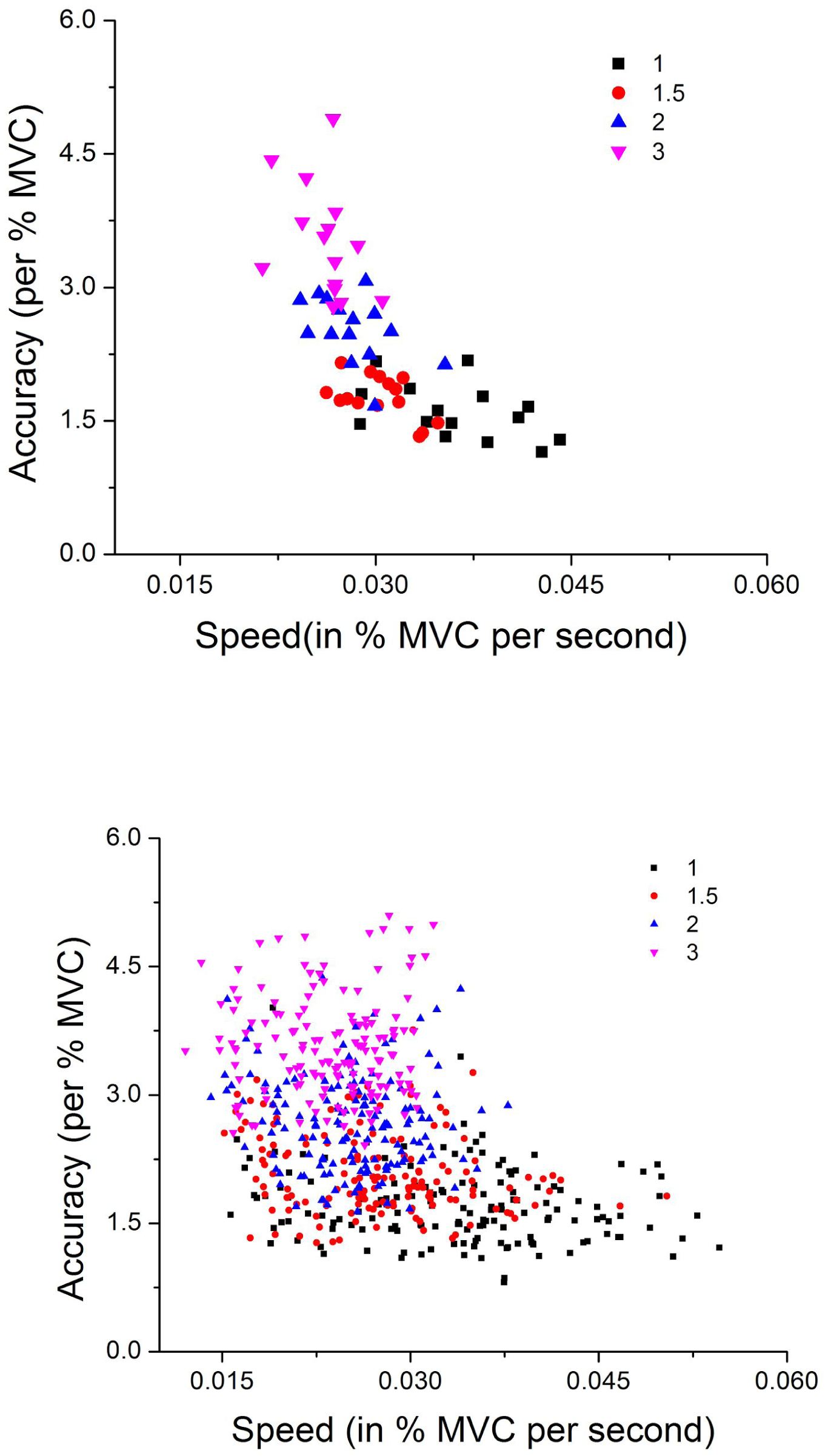
Figure Speed vs accuracy: (Top) Representative single subject. (Bottom) For all 10 subjects.

### Manifestations of biasing

An extended conjecture in terms of independence on this result is the relationship between the independence of participating elements and the motor performance. Despite higher mechanical requirements of employing the index finger (the independent) to produce larger absolute force, the movement control system continues to prefer it as against the little finger (the dependent) which could have produce smaller absolute force. Hypothetically, had the system been purely energy conservative system, then the system should have exploited more of little finger and consequently yield little finger biased trajectories. This is a clear manifestation of the system operating under more than a single objective function. And with these results, at least we can speculate that the control system has a preference of elements which are more independent. The improvement of the tracking performance could be due to the system having had used more of the independent element over against its less independent counterparts. To the least, there may be a causal relationship between them. Of course, similar experiments on a systematic and large set of elemental pairs need to be studied to derive into such a cause and effect global relationship.

And lastly, speculating on the neural control of this behaviour, the complementary measure of the biasing value (from the unbiased condition of zero - the state of balanced neural sharing, *Figure Biasing*) could be used as a relative index of neural biasing which should be present at atleast higher levels of the control hierarchy. At least in principle, the method employed here for measuring neural biasing between the participating elements could be designed into a behavioural basis for characterising neuromotor performance across populations of interest.

## Conclusion

Some behavioural features involved in this task of visuomotor tracking in force-force space have been characterised. These results may imply to a nature of the motor control system which prefers higher independence of the participating elements. This may manifests into improvement of tracking accuracy and control with increasing contribution of relative mechanical effort from the independent element. These results provide insights about how the movement control system realises certain perceived and performing behavioural parameters. It has critical implications in how the control and coordination is achieved in the redundant multi-effector system. Moreover, the methodology adopted for showing the biasing of the system towards any of the participating elements may prove to be useful in quantifying the neural biasing between any elemental pairs.

Further attempts to understand the underlying principles and mechanisms involved in this behaviour of finger force generation through modulated online visual feedback may be achieved through experiments with simpler tasks (maybe such as reaching a point or tracing only a straight line in force-force space). Perturbation studies could reveal functional characteristics; in addition to constant modulation, experiments involving proportional, anti-proportional, directional, and stochastic modulation could be designed. Another set of experiments on speed constrained and/or accuracy constrained tasks could also elucidate the behaviour in question. And lastly, the efforts exerted by the participating elements could be explored beyond the reported ranges and spectrum to establish any possible behavioural global relationships.

## Acknowledgements

Industrial Consultancy & Sponsored Research, Indian Institute of Technology Madras, India for new faculty seed grant.

## References

1. Beer, Randall D. “Beyond control: The dynamics of brain-body-environment interaction in motor systems.” Progress in motor control. Springer US, 2009. 7–24.

2. Bernstein, N. A. “The control and regulation of movements.” (1967).

3. Danion, Frederic, et al. “A mode hypothesis for finger interaction during multi-finger force-production tasks.” Biological cybernetics 88.2 (2003): 91–98.

4. Fitts, Paul M., and James R. Peterson. “Information capacity of discrete motor responses.” Journal of experimental psychology 67.2 (1964): 103.

5. Flash, Tamar, and Neville Hogan. “The coordination of arm movements: an experimentally confirmed mathematical model.” Journal of neuroscience 5.7 (1985): 1688–1703.

6. Guigon, Emmanuel, Pierre Baraduc, and Michel Desmurget. “Computational motor control: redundancy and invariance.” Journal of neurophysiology 97.1 (2007): 331–347.

7. Harris, Christopher M., and Daniel M. Wolpert. “Signal-dependent noise determines motor planning.” Nature 394.6695 (1998): 780–784.

8. Johansson, R. S., and G. Westling. “Roles of glabrous skin receptors and sensorimotor memory in automatic control of precision grip when lifting rougher or more slippery objects.” Experimental brain research 56.3 (1984): 550–564.

9. Johansson, Roland S., and Kelly J. Cole. “Sensory-motor coordination during grasping and manipulative actions.” Current opinion in neurobiology 2.6 (1992): 815–823.

10. Latash, Mark L. “The bliss (not the problem) of motor abundance (not redundancy).” Experimental brain research 217.1 (2012): 1–5.

11. Matthews, P. B. C. “Muscle spindles and their motor control.” Physiological Reviews 44.2(1964): 219–288.

12. Miall, R. Christopher, D. J. Weir, and J. F. Stein. “Intermittency in human manual tracking tasks.” Journal of motor behavior 25.1 (1993): 53–63.

13. O’Sullivan I, Burdet E, Diedrichsen J (2009) Dissociating Variability and Effort as Determinants of Coordination. PLoS Comput Biol 5(4): e1000345. doi: 10.1371/journal.pcbi.1000345

14. Reis, Janine, et al. “Noninvasive cortical Stimulation enhances motor skill acquisition over multiple days through an effect on consolidation.” Proceedings of the National Academy of Sciences 106.5 (2009): 1590–1595.

15. Rothwell, J. C., et al. “Manual motor performance in a deafferented man.” Brain 105.3 (1982): 515–542.

16. Sanes, Jerome N., et al. “Motor deficits in patients with large-fiber sensory neuropathy.” Proceedings of the National Academy of Sciences 81.3 (1984): 979–982.

17. Shmuelof, Lior, John W. Krakauer, and Pietro Mazzoni. “How is a motor skill learned? Change and invariance at the levels of task success and trajectory control.” Journal of neurophysiology 108.2 (2012): 578–594.

18. Slifkin, Andrew B., David E. Vaillancourt, and Karl M. Newell. “Intermittency in the control of continuous force production.” Journal of Neurophysiology 84.4 (2000): 1708–1718.

19. Todorov, Emanuel, and Michael I. Jordan. “Optimal feedback control as a theory of motor coordination.” Nature neuroscience 5.11 (2002): 1226–1235.

20. Vaillancourt, David E., and Daniel M. Russell. “Temporal capacity of short-term visuomotor memory in continuous force production.” Experimental Brain Research 145.3 (2002): 275–285.

21. van Beers, Robert J., Eli Brenner, and Jeroen BJ Smeets. “Random walk of motor planning in task-irrelevant dimensions.” Journal of neurophysiology 109.4 (2013): 969–977.

22. van Duinen, Hiske, and Simon C. Gandevia. “Constraints for control of the human hand.” The Journal of physiology 589.23 (2011): 5583–5593.

23. Wolpert, Daniel M., and Michael S. Landy. “Motor control is decision-making.” Current opinion in neurobiology 22.6 (2012): 996–1003.

24. Zatsiorsky, Vladimir M., Zong-Ming Li, and Mark L. Latash. “Coordinated force production in multi-finger tasks: finger interaction and neural network modeling.” Biological cybernetics 79.2 (1998): 139–150.

